## Supplementary material for "The first complete 3D reconstruction and morphofunctional mapping of an insect eye": Appedix 1

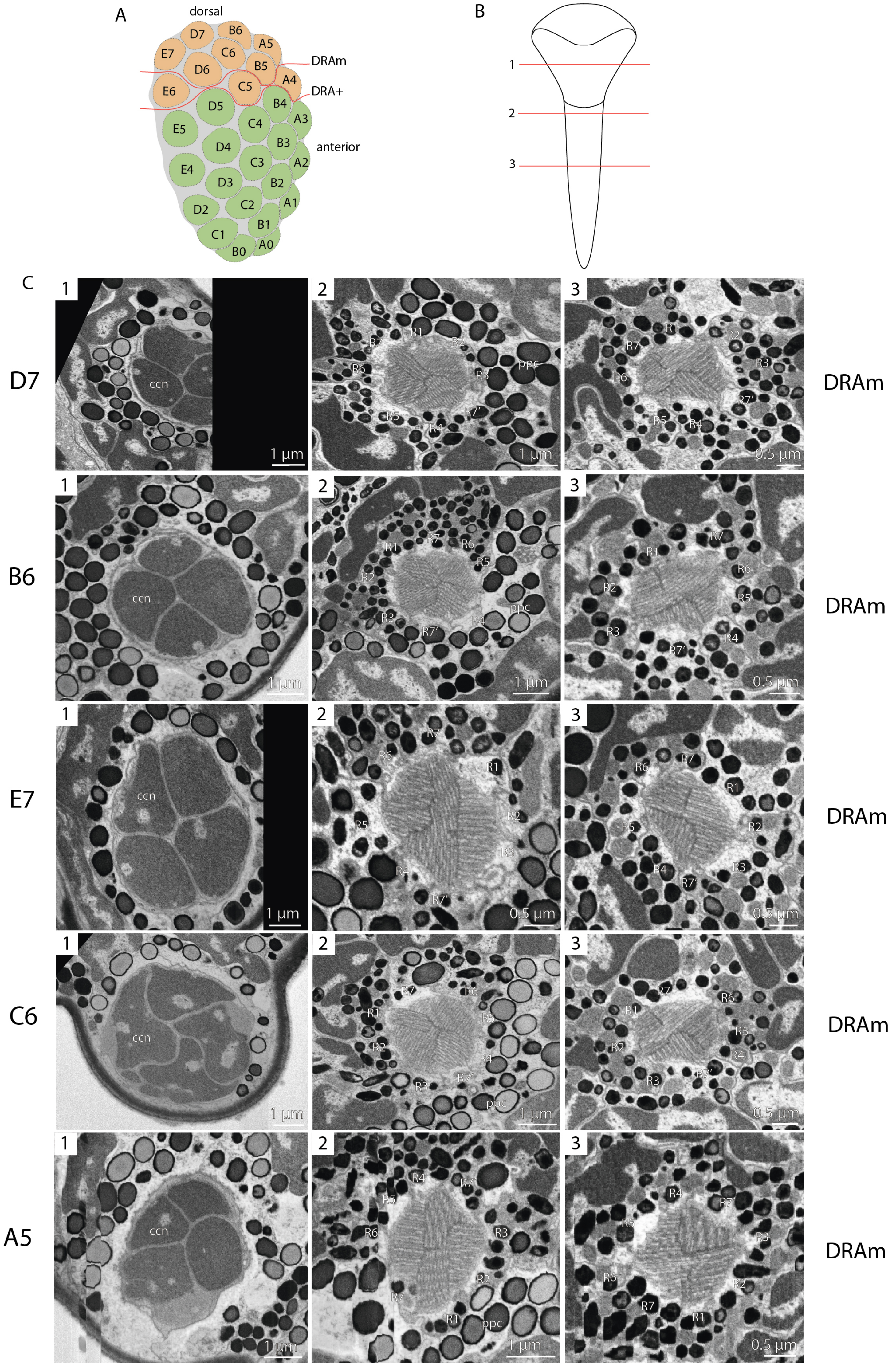

A
dorsal
D7
B6
A5
E7
C6
D6
B5
DRAm
C5
A4
DRA+
E6
D5
B4
A3
C4
E5
B3
D4
anterior
C3
A2
E4
B2
D3
A1
C2
D2
B1
C1
A0
B0
B
1
2
3
C
D7
B6
E7
C6
A5
DRAm
DRAm
DRAm
DRAm
DRAm
ccn
ppc
R1
R2
R3
R4
R5
R6
R7
R7'
1 µm
0.5 µm

Supplement 1C, continuation

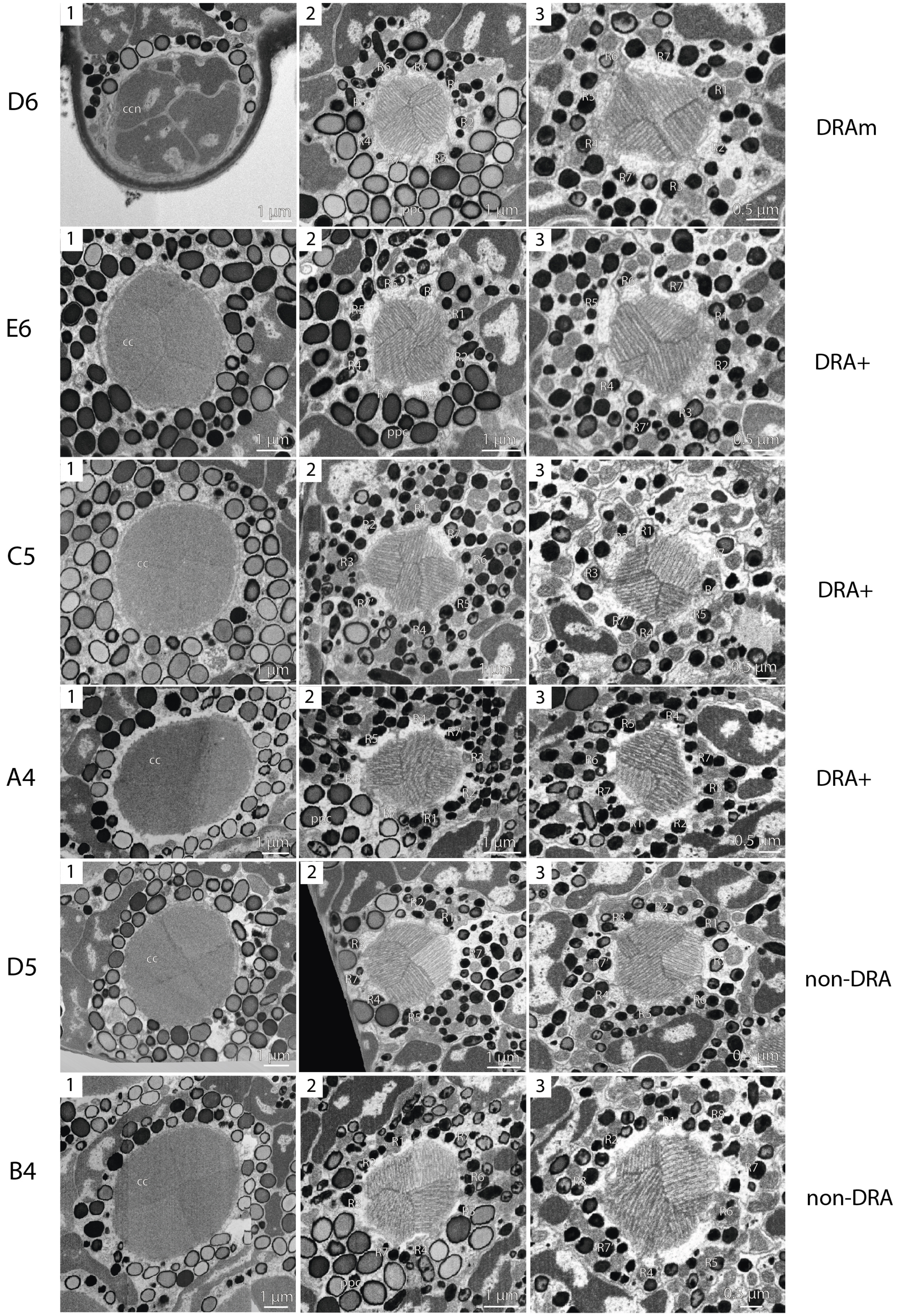

Supplement 1C, continuation

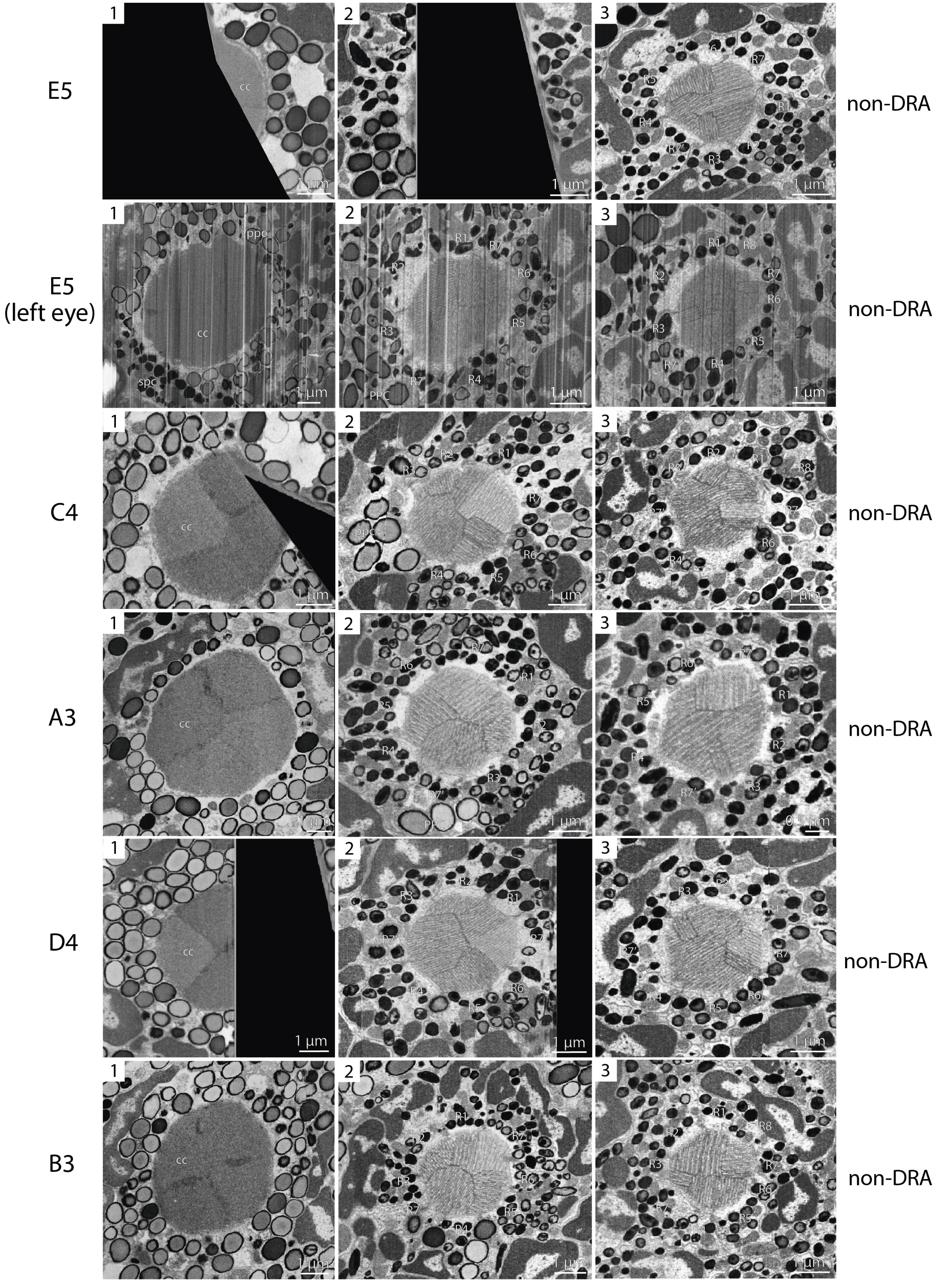

Supplement 1C, continuation

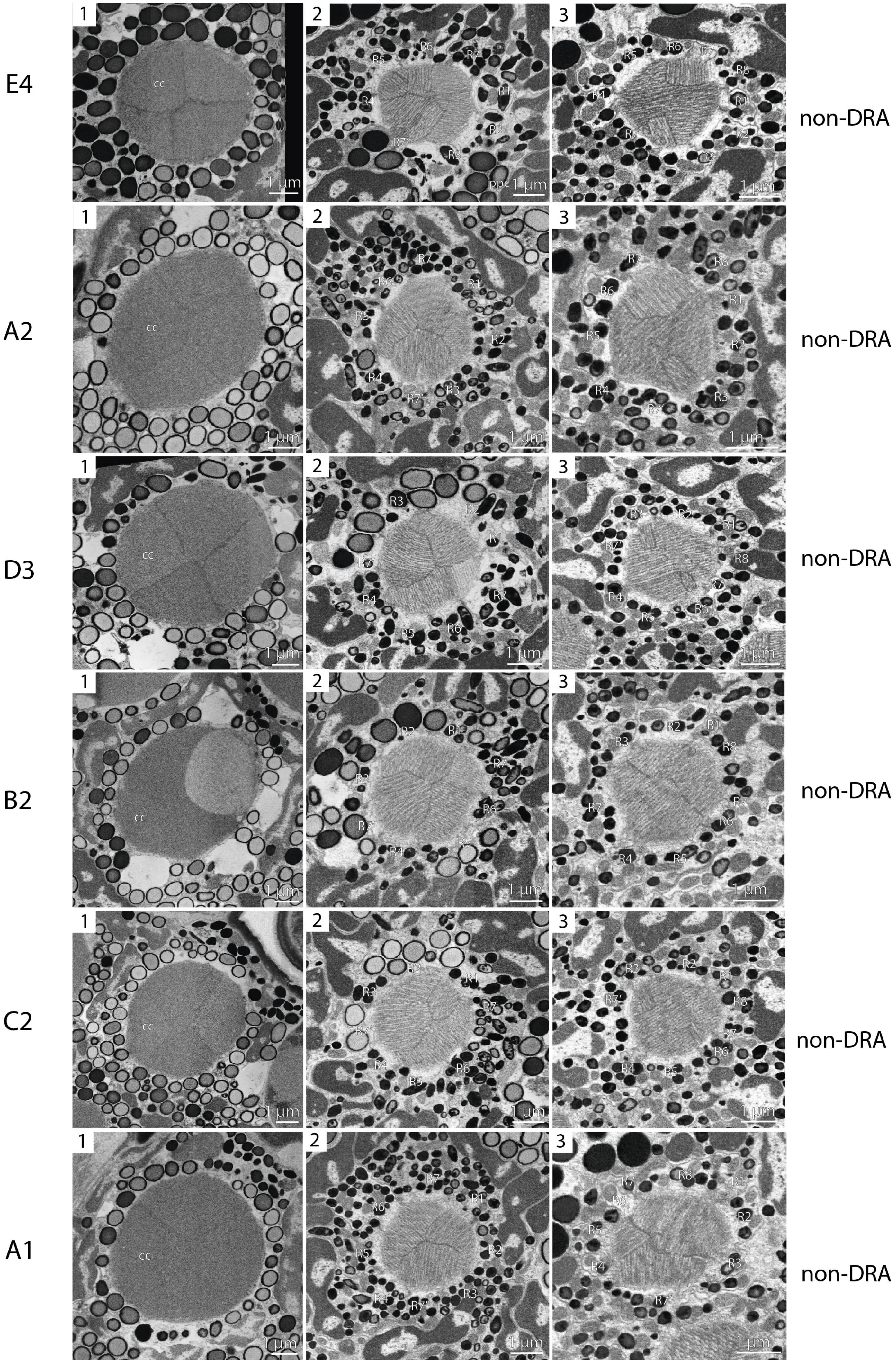

Supplement 1C, continuation

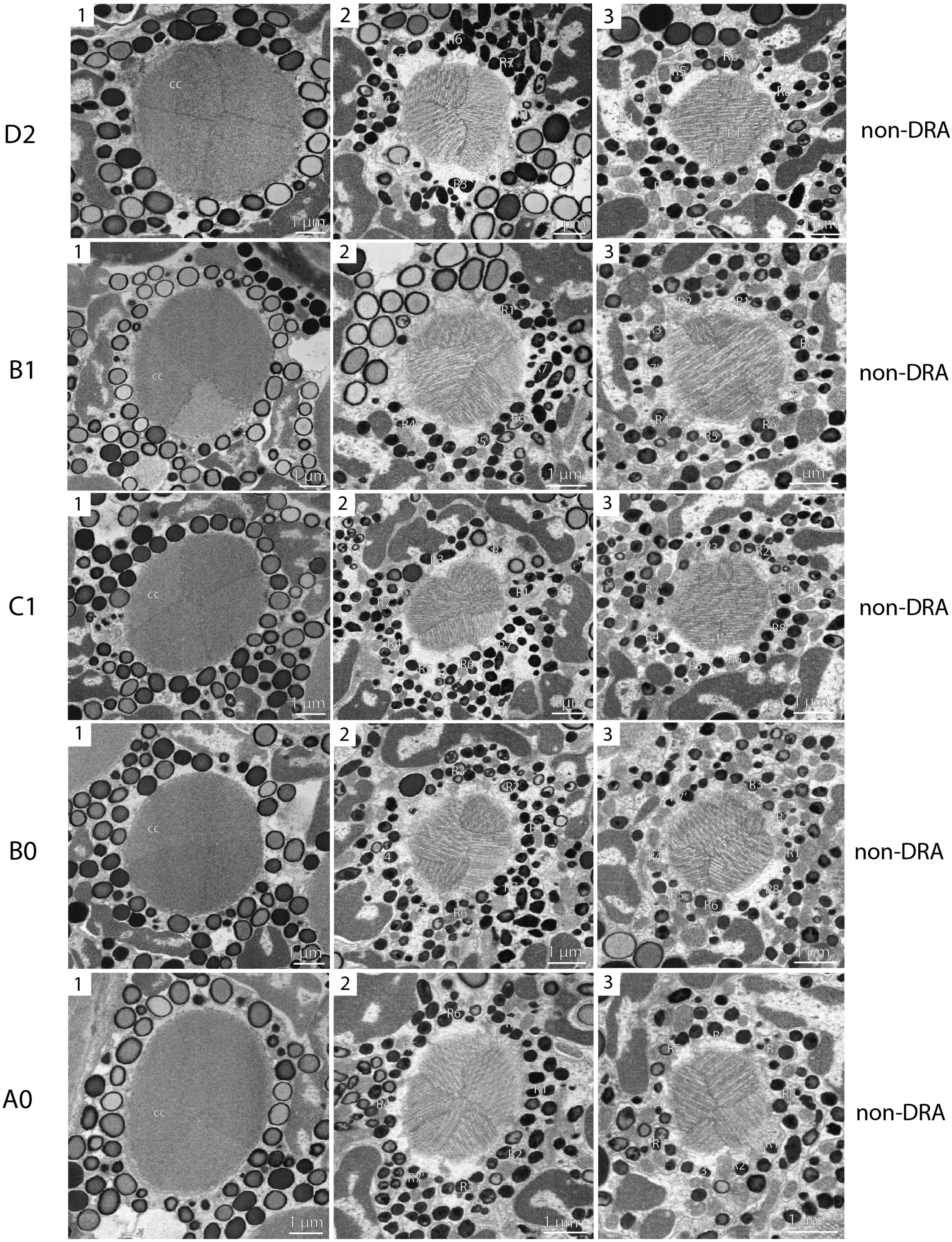
